# Binocular vision emerges from the coordinated development of orbit convergence, eye orientation, and high-acuity retinal specializations

**DOI:** 10.64898/2025.12.19.695568

**Authors:** Alfonso Deichler, Macarena Ruiz-Flores, Natalia I. Márquez, Cristian Morales, Luciana Lopez-Jury, Tomas Vega-Zuniga, Jorge Mpodozis, Macarena Faunes, Gonzalo J. Marín

## Abstract

Binocular vision requires both eyes to be aligned such that their visual fields overlap. A long-standing premise derived from comparative studies is that the orientation of the orbits determines eye position, and thereby the extension of this overlap, the binocular field. In addition, to produce an accurate neural representation, the binocular field must integrate with the position of retinal high-acuity areas and with the extent of uncrossed retinal projections. It remains unknown, however, whether the binocular field is already formed at the time of eye-opening, as well as when and how it integrates with neuroanatomical visual traits during development. Using the diurnal rodent *Octodon degus*, a suitable animal model for visual neuroscience, we combined CT-based 3D cranial reconstructions, quantitative measurements of visual-field geometry, whole-mount retinal topography, neural tracing of retinal projections, and behavioral assays to reconstruct the postnatal assembly of the binocular visual system. We show that orbital and ocular orientations shift substantially after birth, broadening the dorsal binocular field; that retinal ganglion cell distributions sharpen into a horizontal visual streak and a defined *area centralis*; and that ipsilateral projections to the superior colliculus mature in parallel to binocular expansion. These changes coincide with the emergence of binocular-dependent behaviors such as depth discrimination and looming-evoked escape responses. Together, our findings demonstrate that binocular vision emerges through the coordinated alignment of multiple developmental processes across levels of organization.

## Introduction

Binocular vision is the visual perception that occurs with both eyes working simultaneously. It facilitates depth perception, enhances visual acuity, and maximizes sensitivity in low-light environments^1–4^. Consequently, visually active nocturnal animals tend to be more binocular than their diurnal relatives, and hunters are more binocular than prey species^5–8^. The size of the binocular visual field, the portion of space viewed by both eyes, correlates with the convergence of the orbits (eye sockets), thus, it has been proposed that the evolution of forward facing orbits enhances binocularity by positioning the eyes in a corresponding convergent orientation^1,5,9^. Comparative evidence also reveals a consistent correlation between orbit convergence and the location of high-acuity areas in the retina^10^, which determines the portion of the visual field that is scanned at maximal resolution^11,12^. For instance, in mammals with laterally positioned eyes, high-acuity areas—such as the *area centralis* and *fovea*—are confined to the peripheral temporal retina. Species whose eyes face forward, instead, develop specialized areas closer to the retinal center^10–12^, conserving the forward projection of the visual axis (the line of sight to the high-acuity retinal area)^1^. The representation of the expanded binocular field in the brain of frontal-eyed animals is achieved by a proportional increment in the number of retinal axons that do not cross the optic chiasm during development, providing the anatomical substrate for integrating physiological activity of corresponding retinal locations from both eyes^13,14^ (Figure 1a). Therefore, a critical step to achieve binocular perception during development is to align the position of the eyes with multiple other craniofacial and neuroanatomical traits.

**Figure 1.**
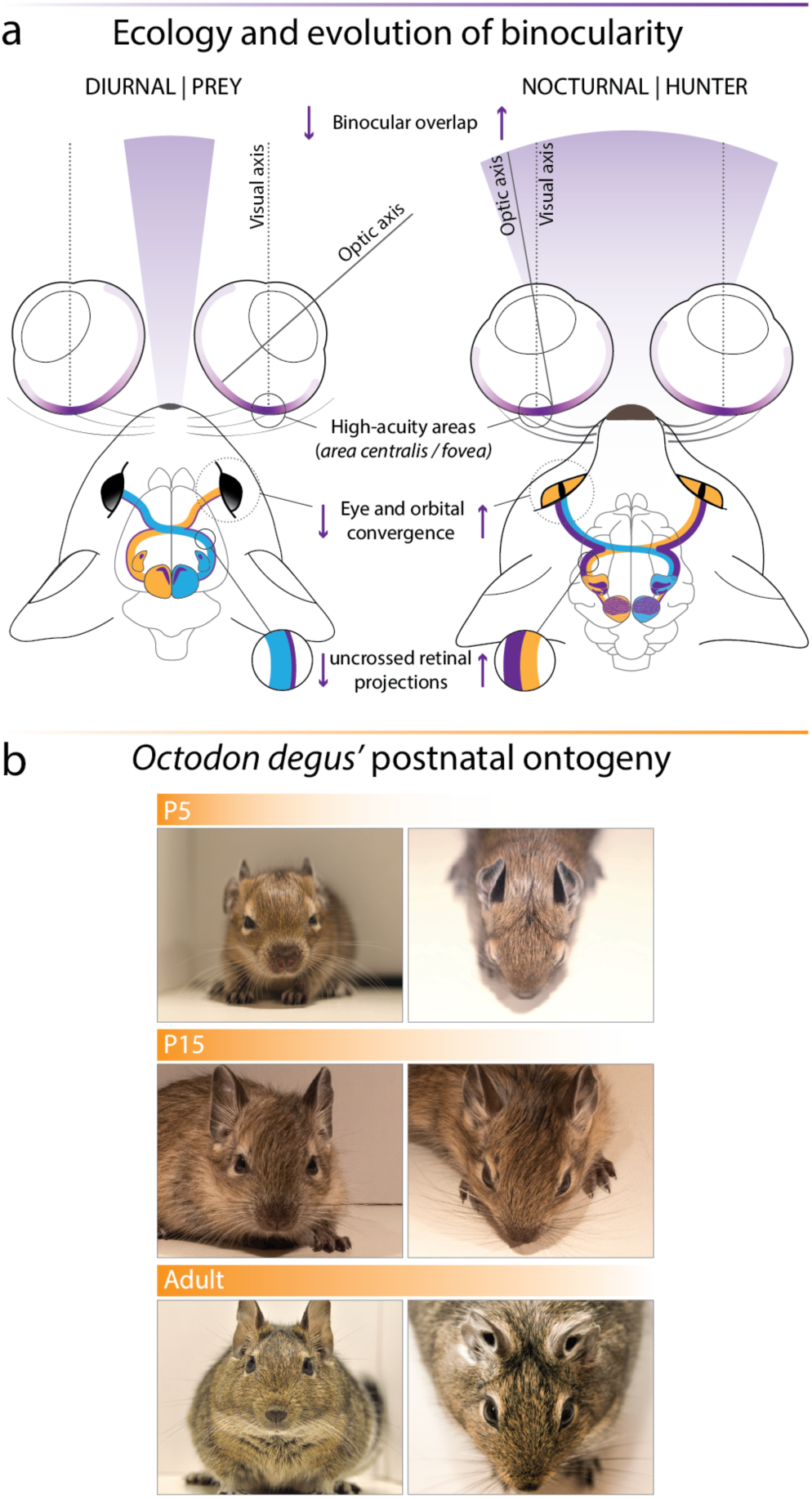
Ecology and ontogeny of binocularity. (a) Conceptual schematics contrasting the integrated phenotypes of animals with laterally placed eyes (diurnal, prey-like) versus convergent eyes (nocturnal, hunter-like). Increasing eye and orbital convergence widens the binocular overlap, brings the visual and optic axes into closer alignment, shifts high-acuity retinal specializations (*area centralis*/*fovea*) toward the retinal center, and increases the proportion of uncrossed (ipsilateral) retinal projections from the temporal retina. (b) Representative photographs of *Octodon degus* across postnatal ontogeny (postnatal day (P) 5, P15, and adult), shown in frontal and dorsal views, illustrating craniofacial growth and progressive eye convergence in this precocial rodent.

Postnatal corrections in eye orientation have been reported in several species, including owls^15^, cats^16^, and humans^17^, during critical periods of plasticity in the anatomy and physiology of binocular areas of the visual cortex^18^. At equivalent critical periods, binocular adaptive behaviors, such as escape and hunting also mature^19–21^, suggesting a global and coordinated postnatal rearrangement of the binocular system. However, it remains unknown whether the monocular and binocular visual field layouts are already formed at birth or undergo further refinement. Variations in the orientation of the orbits and eyes, as well as in the size of each monocular visual field, are expected to produce changes in the binocular visual field^1,5,22^. Yet, how orbital orientation affects binocularity during development has not been explored. It has been suggested that proper ocular position is attained by pointing the high-acuity area of both retinas towards the same target^15^, a critical step to achieve binocular fusion, i.e. the combination of two retinal images into a unified percept. However, high-acuity areas gradually differentiate after birth following massive rearrangements in the number and distribution of retinal ganglion cells^23–26^. Therefore, whether the timing of retinal maturation matches the period of interocular alignment must be determined to test the role of retinal specialization in the development of binocularity, which in turn could reveal a complex interplay between the maturation of the visual fields, eye and orbital orientation, retinal topography, and behavior.

To determine when and how the different components of the binocular system become functionally integrated during development, we investigated the postnatal development of binocularity in the highly visual diurnal rodent, *Octodon degus*^27^. This species is born in an advanced precocial state and becomes visually active shortly after birth and provides an excellent model for studying the postnatal development of visual system (Figure 1b). By characterizing the postnatal maturation of visual fields, orbital geometry, retinal topography, central projections, and visually guided behaviors, we show that binocularity is not fully established at eye opening. Instead, multiple anatomical features of the visual system undergo coordinated postnatal changes that parallel the maturation of binocularly guided behaviors. Together, these results frame binocular vision as an emergent functional trait that arises from the coordinated timing of structural and behavioral development.

## Results

### Coordinated development of the binocular field and orbit convergence

To quantify postnatal eye convergence in degus, we measured the extension and orientation of the monocular visual fields and calculated the amount of binocular overlap at different postnatal stages. Degus are particularly well-suited for these procedures, as their eyes are already well-developed and prominent at early postnatal stages (figure 1b), allowing for the mapping of visual fields with a standard ophthalmoscopic reflex technique^7,28–30^. Despite their advanced precociality, we found that the visual field organization is far from mature in newly born degus. Binocular overlap, initially confined to a small part of the visual field, expands steadily and significatively throughout the first postnatal month (ANOVA: *F*_(4,18)_ = 12.96, *p* < 0.001). At postnatal day 5 (P5) the binocular field forms a narrow band with a maximum azimuthal extension of 25.5 ± 5.1° (mean ± SD) that expands to 54.2 ± 6.8° by P30 (figure 2a, b). From P30 to adulthood (individuals older than 5 months), the dorsal portion of the binocular field widens by an additional 5° (figure 2a; supplementary table 1). In a separate cohort of animals on which we measured the projection of the optic axis in the campimeter as a proxy for eye position, we found a progressive elevation from 33.6 ± 12.5° at P5 to 63 ± 5.2° in adults (Kruskal-Wallis: *H* = 18.16, *p* = 0.001; figure 2c; supplementary table 1), a result consistent with the convergence of the eyes in an upward orientation. In addition, monocular fields also broaden (figure 2d) from an average 166.6 ± 15.3° in P5 to 186.3 ± 7.2° in adults (Kruskal-Wallis: *H* = 16.194, *p* = 0.003; supplementary table 1). These results show that monocular fields expand and, simultaneously, converge due to the upward and medial reorientation of the eyes, resulting in an enlarged binocular overlap.

**Figure 2.**
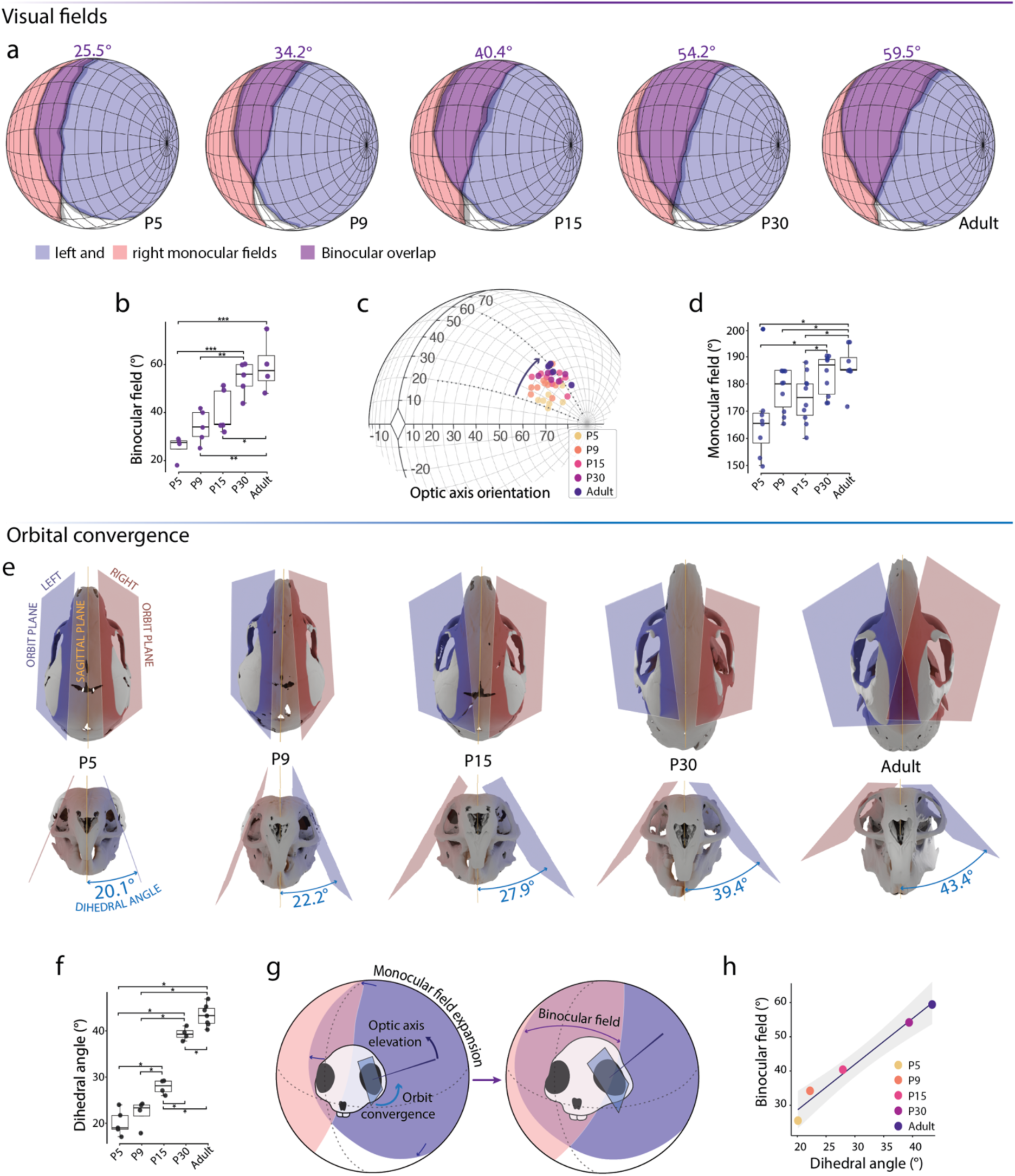
Postnatal development of binocular craniofacial anatomy in *Octodon degus*. (a) Orthographic projections of the mean visual fields upon a sphere centered on the animal’s head at each developmental stage. Mean maximal azimuthal extension of the binocular visual field is indicated for each stage. Shaded area at the border of each monocular field represents ± s.e.m. Sample sizes: P5, *n*=4, P9, *n*=5, P15, *n*=5, P30, *n*=5, and adult, *n*=4. (b) Box plots displaying maximum binocular overlap across postnatal stages. (c) Orthographic projection of the optic axis at the evaluated developmental stages. Measurements were taken in both eyes but are shown on the left side for clarity. (d) Box plots displaying the monocular field extent across postnatal stages. (e) Schematics depicting the orbital planes at each stage in dorsal (top row) and frontal (bottom row) views of representative CT-scanned crania. Mean dihedral angle values for each stage are indicated. Note that larger dihedral angles indicate more convergent orbits. Sample sizes: P5, *n*=5; P9, *n*=4; P15, *n*=4; P30, *n*=4; and adult, *n*=7. (f) Box plots displaying orbital convergence (dihedral angle) across postnatal stages. (g) Schematics summarizing the correlation between the expansion and overlap of the monocular visual fields, the elevation of the optic axes, and the convergence of the orbits. (h) Ontogenetic relationship between orbital convergence and binocular field. Points are mean values for each stage. The line shows Pearson’s correlation with 95% confidence bands in gray. The linear regression is significant (binocular field vs dihedral angle:(r = 0.99, 95%, CI = 0.82–1.00, n = 4, t(3) = 10.93, p = 0.0016). Statistically significant differences determined using paired samples t-tests (b), and Wilcoxon rank-sum tests (d and f): \**p* < 0.05; \*\**p* < 0.01; \*\*\**p* < 0.001.

Reorientation of the optic axis may result from changes in eye position within the orbits, a reorientation of the orbits themselves or a combination of both processes. To test these possibilities, we quantified postnatal changes in orbit orientation. Using 3D rendered models of CT-scanned skulls, we calculated the dihedral angle formed by the intersection between the orbital margin plane and the midsagittal plane as a measure of orbit convergence (where larger dihedral angle values indicate more convergent orbits)^9^. As observed with the visual field, orbit morphology is still immature shortly after birth. At P5 the orbits appear relatively flat from a dorsal view (top row in figure 2e) with a mean dihedral angle of 20.1 ± 2.7° (figure 2e, f). This angle increases steadily, and in a statistically significant manner, thereafter to an average of 43.4 ± 2.3° in adults (Kruskal-Wallis: *H* = 20.972, *p* < 0.001; supplementary table 1), producing upward-converging orbits (figure 2e, f). These data indicate that the orbital plane rotates upwards in concert with the eyes, a progression that is consistent with the upward facing of the eyes (Figure 1b) and the dorsal expansion of the binocular field. Consequently, the correlation between orbit convergence and binocular overlap previously established at the phylogenetic level^9^ is recapitulated during postnatal ontogeny in the degus (figure 2g, h).

### Retinal topography develops in parallel to the visual field layout

In degus, retinal topography is coherent with visual field extension and orientation, following general patterns also found in other rodent species^7^. The peak of retinal ganglion cell (RGC) density is located in the dorsotemporal retina (shown in a representative adult retinal map in the right corner of figure 3a), forming the *area centralis* that points to the frontal inferior binocular field^7^. High-RGC density extends further nasally from the *area centralis,* forming a band or “visual streak” above the retinal equator, which enables high-acuity panoramic vision across the visual horizon in preyed species inhabiting open grasslands, such as the degus^6,10^. RGC density decreases more sharply dorsal to the streak than ventral to it, resulting in a relatively denser ventral retina, which may enhance sensitivity to aerial predators appearing in the dorsal binocular field^11,28,31^.

**Figure 3.**
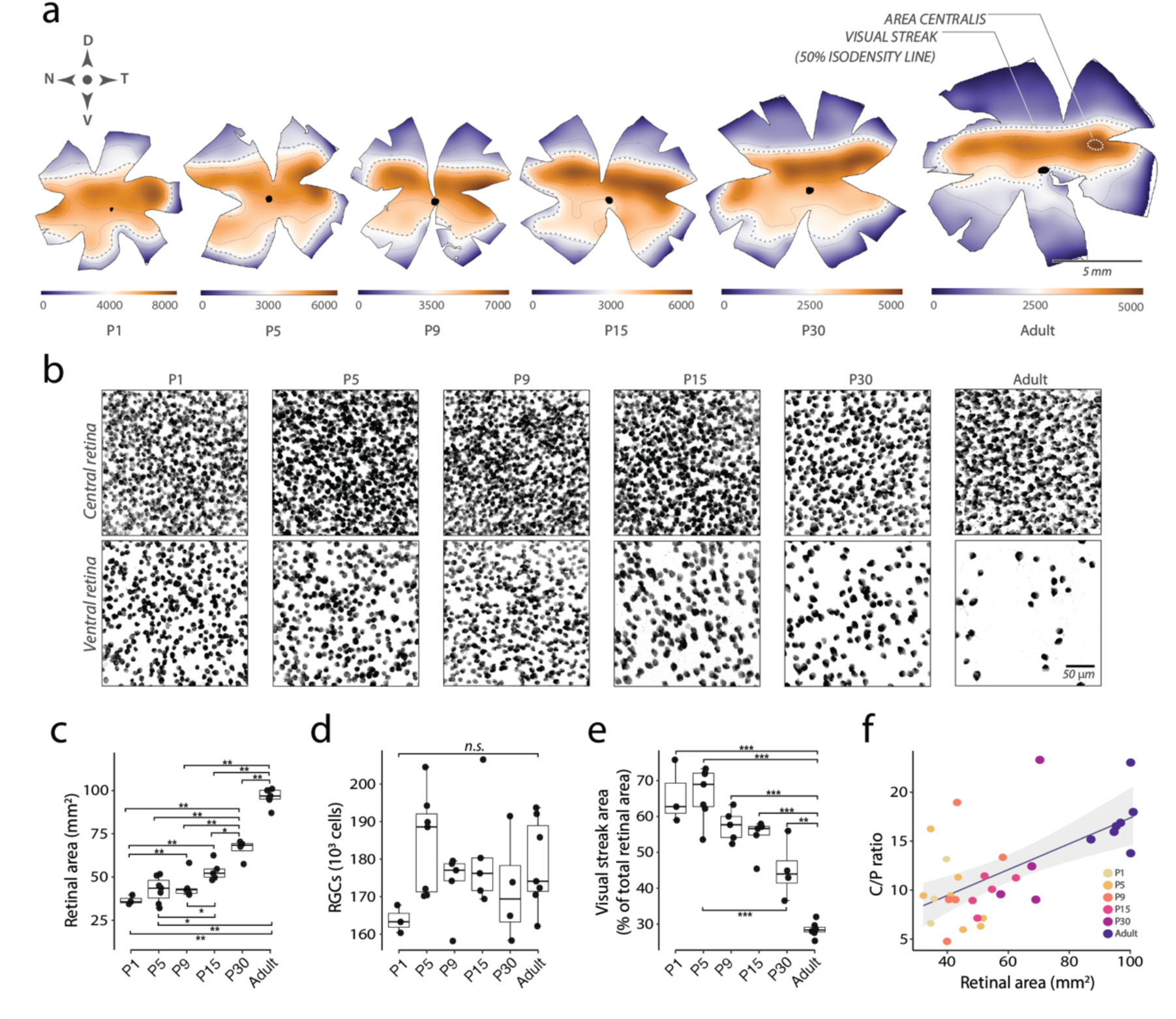
Postnatal development of retinal specialization. (a) Isodensity maps of labeled RGCs on flat-mounted retinas (one representative case per stage). Warm colors indicate higher densities; the 50% isodensity contour (dotted gray line) delineates the visual streak and the dotted white line marks the area centralis. Sample sizes: P1, *n*=3; P5, *n*=7; P9, *n*=5; P15, *n*=5; P30, *n*=4; adult, *n*=7. (b) Photomicrographs of RBPMS immuno-stained RGCs from central and ventral retina across postnatal development. (c – e) Box plots across postnatal stages of (c) retinal area, (d) total RGC number, and (e) the proportion of retina occupied by the visual streak (region enclosed by the 50% isodensity contour). (f) Relationship between C/P ratio (central-to-peripheral RGC density ratio) and retinal area. Linear regression analysis shows that retinal area significantly predicts C/P (β = 0.13, SE = 0.03, t(29) = 4.29, p < 0.001), accounting for 39% of the variance (R² = 0.39, *F*_(1,29)_ = 18.44, *p* < 0.001); points are individual retinas, colored by stage. Pairwise comparisons using Wilcoxon rank-sum test: \**p* < 0.05; \*\**p* < 0.01; \*\*\**p* < 0.001. *n.s.*, not significant.

Based on experiments inducing strabismus (eye misalignment) by rearing animals with one eye occluded, it has been proposed that interocular alignment is achieved by the binocular conjugation of the *area centralis* (i.e. by coordinately pointing the areas from both eyes toward the same location in space)^15^; however, this would require these high-acuity areas to be formed before (or at least during) eye alignment, a condition that remains untested. To address this hypothesis, we examined the maturation of the retinal mosaic. Flat-mounted retinas were immunolabelled with the pan-RGC marker Rbpms^32^, and isodensity maps were constructed across postnatal stages. In early stages, we found the peak RGC density occupies a broad horizontal region of the retina. By P15 the visual streak is discernible, though still not completely differentiated from the ventral retina (figure 3a, b). Even at P30, the distinction between the visual streak and a discrete *area centralis* remains less pronounced than in the adult (figure 3a).

Next, we explored the process that generates the local differences in cellular density. Previous studies in rabbits, cats, and other mammals have shown that non-uniform retinal growth is the main driver of the emergence of retinal topography: once the total number of retinal ganglion cells (RGCs) reaches a plateau, the peripheral retina continues to expand faster than the center, producing a steeper decline in cell density toward the edges^23,24,33–35^. Consistent with these studies, our data in degus (summarized in supplementary table 2) show that, while the total number of RGCs stabilize and show no further statistical changes after P5 (ANOVA, *F_(5, 25)_*, *p* = 0.229), the retina continues to grow significatively at least until after P30 (Kruskal-Wallis: *H* = 28.754, *p* < 0.001; figure 3c, d). Meanwhile, we found that density gradients sharpen: the high-density band demarcated by the 50 % isodensity contour—destined to become the adult visual streak—shrink from ∼60–70 % of retinal area at P1 to ∼20 % in adults (Kruskal-Wallis: *H* = 24.118, *p* < 0.001; figure 3a, e). Additionally, the central-to-peripheral (C/P) density ratio increases with retinal size (linear regression: slope = 0.126 ± 0.028, *p* < 0.001), rising from approximately 9:1 at P1 to around 16:1 in adults (figure 3b, f). Together, these findings support a model in which asymmetric retinal growth shapes retinal specializations as the binocular visual field expands.

### The development of uncrossed retinal projections predates the postnatal maturation of the binocular field and retinal topography

The distribution of RGCs projecting to the ipsilateral brain matches the projection of the binocular field on the retina. Thus, the number of retinal axons that do not cross at the optic chiasm is proportional to the degree of binocularity^9,13,36^. In all mammals described so far, the arrival and eye-specific segregation of retinal axons that define binocular territories in the superior colliculus (SC) and the dorsal lateral geniculate nucleus (dLGN), the two principal retinorecipient nuclei, are established before eye opening^37–41^, following a conserved developmental chronogram^42^. However, given that both the extent of binocular overlap and the retinal topography change dramatically after eye opening in the degus, we sought to determine whether ipsilateral projections undergo postnatal maturation by injecting a neural tracer (B subunit of cholera toxin, CTB) into one eye at different postnatal stages (figure 4a).

**Figure 4.**
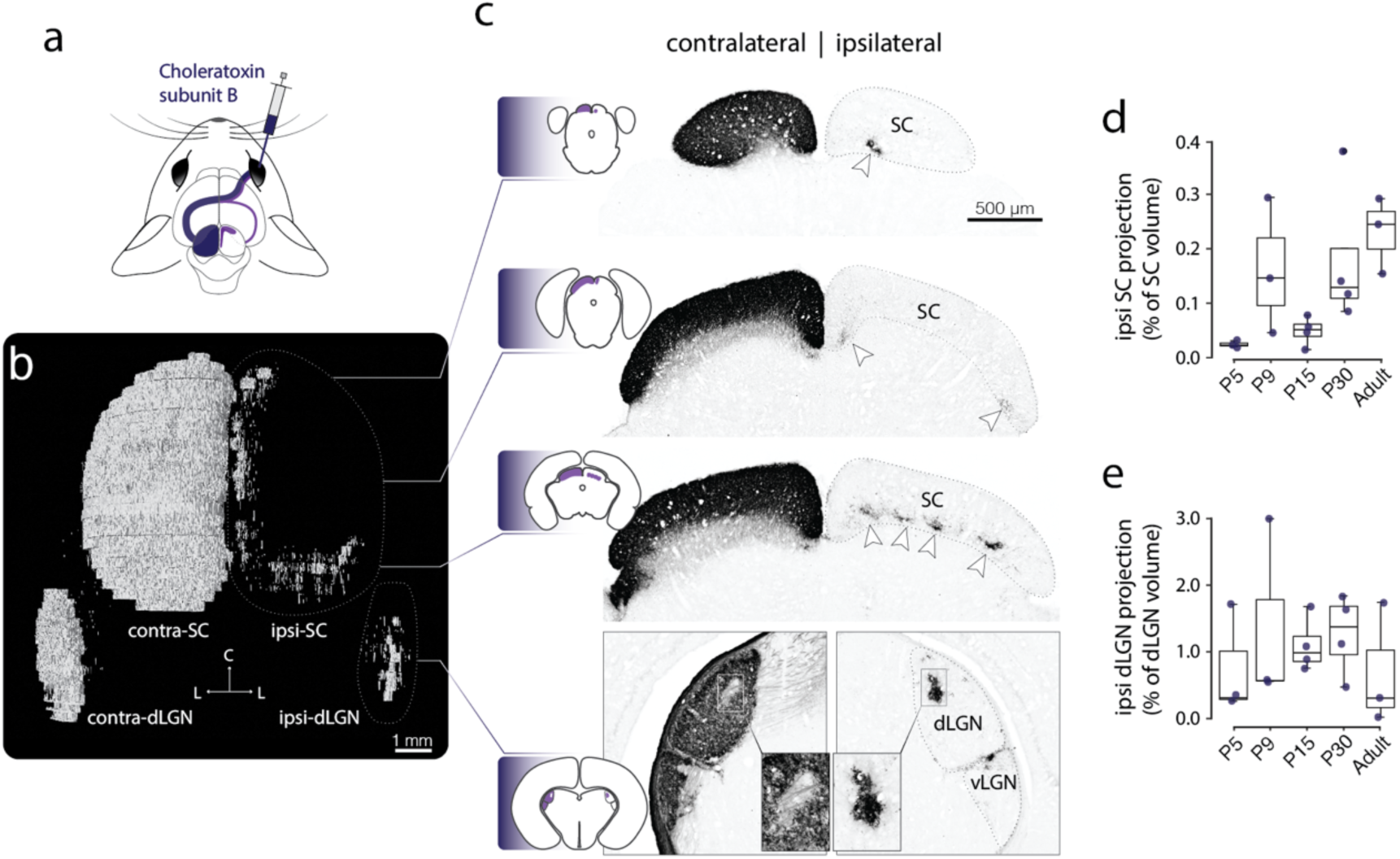
Binocular retinorecipient territories are established before interocular alignment and retinal map refinement. (a) Schematic of the experiments. Intraocular injections of the neural tracer choleratoxin subunit B label retinal fibers and terminals in the ipsilateral and contralateral midbrain and thalamus. (b) Dorsal view of a 3D reconstruction of aligned sections of a representative case from an individual injected at postnatal day 4 and collected at P5. (c) Photomicrographs of selected sections from the case depicted in (b), showing CTB labeling in the superior colliculus (SC) and dorsal lateral geniculate nucleus (dLGN). Ipsilateral projections form a patchwork of terminals all along the rostral and medial SC (arrowheads); in the dLGN ipsilateral terminals occupy a dorsomedial patch that aligns with the contralateral band lacking terminals (bottom insets). (d – f) Box plots representing the relative volume of ipsilateral retinal projections across postnatal stages in the SC (d), and dLGN. Sample sizes are: P5, *n*=3; P9, *n*=3; P15, *n*=4; P30, *n*=4; adult, *n*=3.

Consistent with the previous reports, we found that by P5, the overall retinal projection pattern already mirrors that described for adult degus^7^ and other rodents^28,43–46^ (figure 4b): Contralateral retinal axons densely occupy the superficial layers of SC, whereas ipsilateral input is mostly confined to the deeper optic layer, forming a rostral patchy array that further extends medially and caudally consistent with the collicular representation of the binocular field in rodents^27,47–49^(figure 4b, c). In dLGN, contralateral fibers covered almost the entire nucleus, sparing only a narrow, band-shaped gap in the dorsal medial aspect of the nucleus. Ipsilateral terminals, in turn, form a single, well-defined band of terminals that matches the contralateral gap (figure 4b, c), indicating that eye-specific segregation is already established at this stage.

Although an adult-like projection pattern is already evident by P5, the quantitative analysis (summarized in supplementary table 3) of the relative volume occupied by ipsilateral terminals in the SC revealed a progressive postnatal expansion (Kruskal–Wallis: *H* = 10.96, *p* = 0.026; Jonckheere–Terpstra: *p* = 0.0014; figure 4d). By contrast, the relative ipsilateral input to the dLGN remained statistically unchanged across development (ANOVA, *F*_(4,12)_ = 0.401, *p* = 0.804), occupying ∼1.1 ± 0.8% of the nucleus at all stages (Figure 4e).

These results indicate that the segregation of binocular territories in the SC and dLGN is established before interocular alignment and the expansion of binocular overlap, and may shape the degree of alignment needed to elicit binocular activation of postsynaptic responses in central visual circuits. Moreover, although SC projections display an adult-like pattern by P5, our volumetric analyses show that ipsilateral inputs continue to refine during the period in which binocular alignment and retinal topography mature. Together, the set anatomical changes described so far likely provide the developmental context within which early binocular behaviors begin to emerge.

### Visually guided behaviors emerge during the period of anatomical maturation

To determine whether postnatal changes in the binocular anatomy are related to the maturation of visually guided behaviour, we used two assays of binocular function: the visual cliff test^50,51^, and the looming response test^52,53^.

The visual cliff test measures depth perception, which depends on binocular clues^51,54^. In this test, we placed the animals on a horizontal glass platform that spanned a “shallow” side and a “deep” side, created by positioning patterned surfaces at different depths beneath the glass. We then evaluated their behavior as they approached the boundary between the shallow and deep regions. Notably, we found a significant association between age and depth perception (Fisher’s exact test, *p* <0.001). At early stages (P5 and P9), when binocular overlap is considerably narrower, every animal tested crossed the cliff border (5 out of 5 animals at P5; 6 out of 6 animals at P9). A clear shift in behavior occurred from P15 onwards, however, with most animals stopping at the edge upon their first encounter with the cliff (5 out of 6 animals at P15; 7 out of 8 adult animals) (figure 5b; Supplementary video 1). These results indicate that depth discrimination develops in degus between P9 and P15.

**Figure 5.**
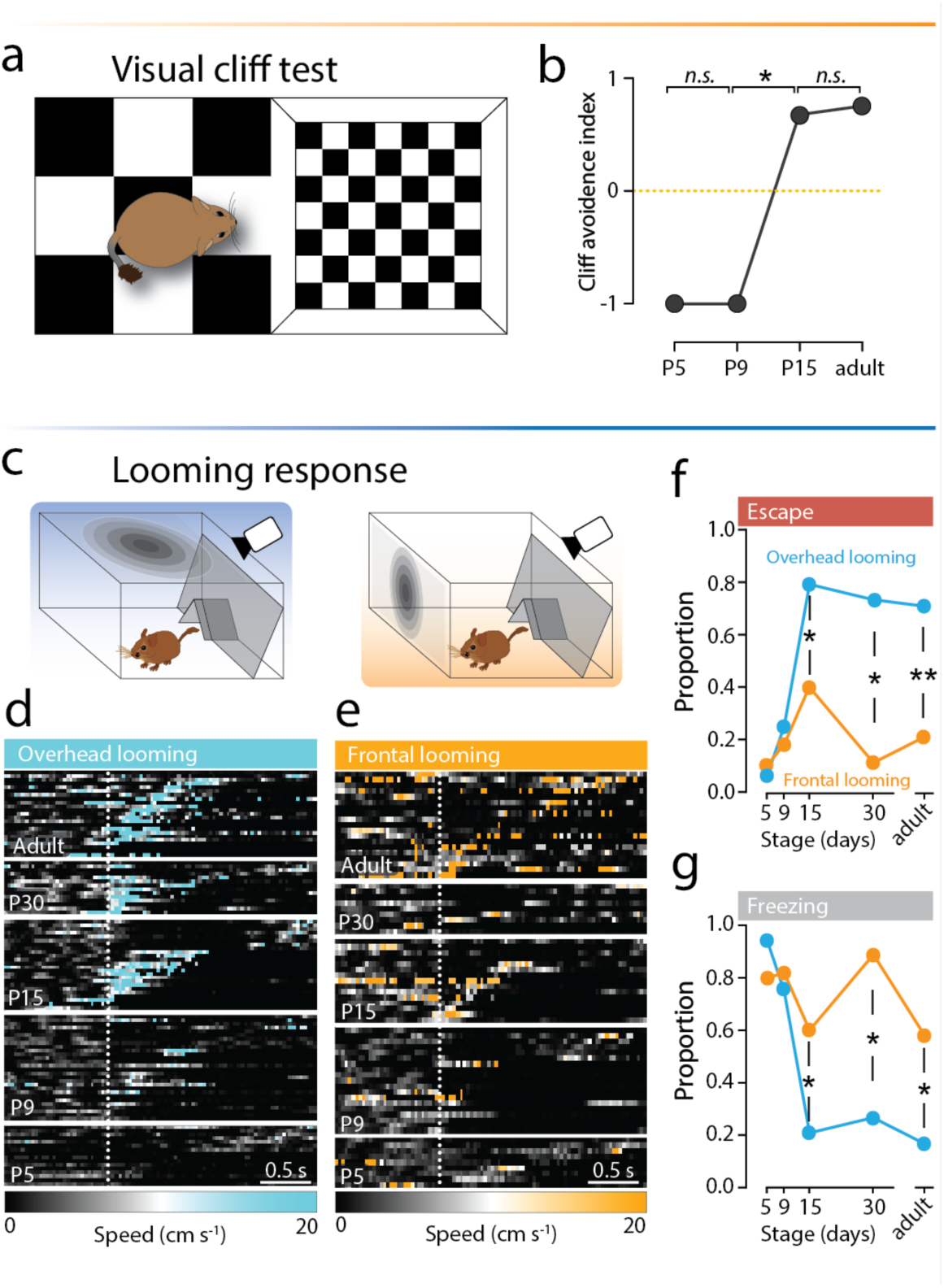
Postnatal maturation of visually guided behaviors. (a) Schematic of the visual cliff setup and (b) Cliff-avoidance index across stages; dashed yellow line marks 50% chance. Each animal underwent one trial at only one stage; sample sizes are: P5, *n*=5; P9, *n*=6; P15, *n*=6; adult, *n*=8. (c - g) Looming response test. Schematics of the looming setups showing overhead at the left and frontal looming at the right. (c) Raster plots of instantaneous speed (cm·s⁻¹) during overhead (d) and frontal (e) looming; rows are trials grouped by stage. Vertical dashed lines indicate stimulus onset (t = 0); scale bar, 0.5 s. (f, g) Proportion of trials showing escape (e) or freezing (f) at each stage for overhead (blue) and frontal (orange) looming. Pairwise comparisons using Fisher’s exact test: \**P* < 0.05; \*\**P* < 0.01. *n.s.* = not significant.

Visually evoked defensive responses also depend on the integrity of binocular connections. Both monocular enucleation^55^ and aberrant ipsilateral retinal projections in mutant mice^56^ reduce escape probability. Consistently, escape reactions to visual threats are stronger in response to stimulation directed to the overhead binocular overlap than to stimulation in other parts of the visual field in degus and other rodents^47,52,57,58^. To determine whether this behavioral asymmetry emerges in concert with binocular development, we exposed individuals across different ages to looming stimulation in the upper and frontal regions of the visual field (figure 5c, d). At early stages (P5 – P9), most individuals indistinctively froze in response to stimulation either in the overhead or frontal visual field (16 out of 17 to overhead looming, 8 out of 10 to frontal looming at P5; 21 out of 28 to overhead looming, 18 out of 22 to frontal looming at P9) (figure 5e, f). From P15 onwards, frontal looming continued to elicit freezing responses in most cases (7 out of 15 at P15; 7 out of 9 at P30; 11 out of 19 adult animals). On the other hand, overhead stimulation responses switched drastically from freezing to escape behavior (19 out of 24 at P15; 11 out of 15 at P30; 11 out of 19 adult animals) (figure 5e, f; Supplementary video 2).

Altogether, these results show that binocular behaviors emerge in concert with the maturation of binocular anatomy. The appearance of depth responses is concurrent with the refinement of high-acuity retinal areas at P15, and the ongoing interocular alignment, whereas the overhead-biased switch from freezing to escape is consistent with the asymmetric expansion of the upper portion of the binocular visual field, and of ipsilateral retinal projections to the SC.

## Discussion

Our results show that binocular vision emerges through a coordinated postnatal alignment of multiple anatomical and functional components of the visual system. In degus, postnatal remodeling of the binocular and monocular visual fields, orbital geometry, and retinal organization occurs in parallel with the maturation of visually guided behaviors that depend on binocular input. These findings emphasize that the development of binocular vision cannot be attributed to a single anatomical change, but instead reflects the integration of sensory pathways, retinal specializations, and craniofacial morphology that together determine how the visual surface is positioned and used in space.

Previous comparative studies in adult mammals have shown a strong correlation between orbital convergence and the maximal extension of the binocular field^9^, however, developmental evidence supporting this observation was previously absent. Our results show that, in degus, the orbital planes converge postnatally while the animal is visually active. The optic axis increases in elevation and the binocular overlap expands, indicating that the eyeball closely follows the reorientation of the orbits. This could be explained by the passive displacement of the eyes driven by the changing orbits. Alternatively, mechanical forces exerted by muscular activity on growing tissues can induce significant morphogenetic changes in the surrounding skeleton. For example, cases involving early eye removal in humans^59^ and laboratory animals (cats^59^, rabbits^60^, lambs^61^) produce arrested orbital development, resulting in a reduced orbital volume and a flattened appearance of the zygomatic arc, reminiscent of the flattened orbits we observed in degus pups. In this scenario, instead of the eye being passively rotated by the orbits, ocular growth, and oculomotor activity associated with exploratory behavior may actively contribute to shape and orient the orbits.

Our data indicate that, in addition to orbital convergence, the expansion of the monocular fields also plays a significant role in the increase of binocular overlap. In his seminal treatise, Walls^5^ pointed out that the extent of the monocular field is an important determinant of binocular overlap, and that it can vary significantly among species. However, subsequent studies using the orbital dihedral angle as a proxy for binocular extension implicitly assumed orbit (eyeball) orientation as the sole determinant of binocularity, overlooking the contribution of monocular field size^62–64^. The primary anatomical factors determining the angular extent of the monocular field include ocular optics and retinal coverage, which is the proportion of the eyeball occupied by the retina^5,22^. In humans, the lens shape flattens postnatally, decreasing its refractive power, which in turn influences the developmental process that brings the mature eye into proper focus^65^. Our results showing substantial postnatal retinal growth are consistent with previous results. In rabbits, for instance, changes in retinal conformation during embryonic and postnatal development result in reduced retinal size relative to eye volume^24^. Whether analogous changes in the optics and retinal coverage occur in other taxa and how these processes integrate with eye growth to influence the size of the monocular visual field are important questions to produce a clearer picture of how the monocular field develops.

Eye growth is also critical for shaping retinal maps, and thus the topography of the monocular images. Here, we show that, although the definitive number of RGCs is already established at early postnatal stages, the topography of their distribution remains immature. This is consistent with the hypothesis of non-uniform growth^23,24,35^, according to which retinal specializations become progressively more defined as the eye enlarges and the retinal surface expands. This refinement of high-acuity areas may be pivotal in guiding ocular orientation adjustments. Experimental induction of strabismus through monocular occlusion in cats^66^, monkeys^67^, and owls^15^ suggests that the conjugated activation of the *area centralis* or *fovea* from both eyes is necessary for postnatal interocular alignment. A well-defined *area centralis* may be required to achieve binocular fusion in the frontal visual field, while refinement of the visual streak may enable precise ocular alignment with the visual horizon, thereby favoring panoramic vision. Both retinal features were immature at early postnatal stages in degus, suggesting that binocular fusion and ocular horizontal alignment proceed while the animal is already able to explore its environment. Furthermore, as retinal specializations become increasingly defined and the monocular image increases in size and acuity, it would be expected that the oculomotor control of gaze orientation and stabilization would become progressively more accurate. It is also expected that by aligning the monocular images from topographically matching areas of the retina, the already established ipsilateral retinal projections would start to generate functional binocular responses in the visual centers of the brain^15^. In this way, ocular and retinal growth are likely integrated with oculo-orbital alignment, oculomotor control, and visual behavior (figure 6).

**Figure 6.**
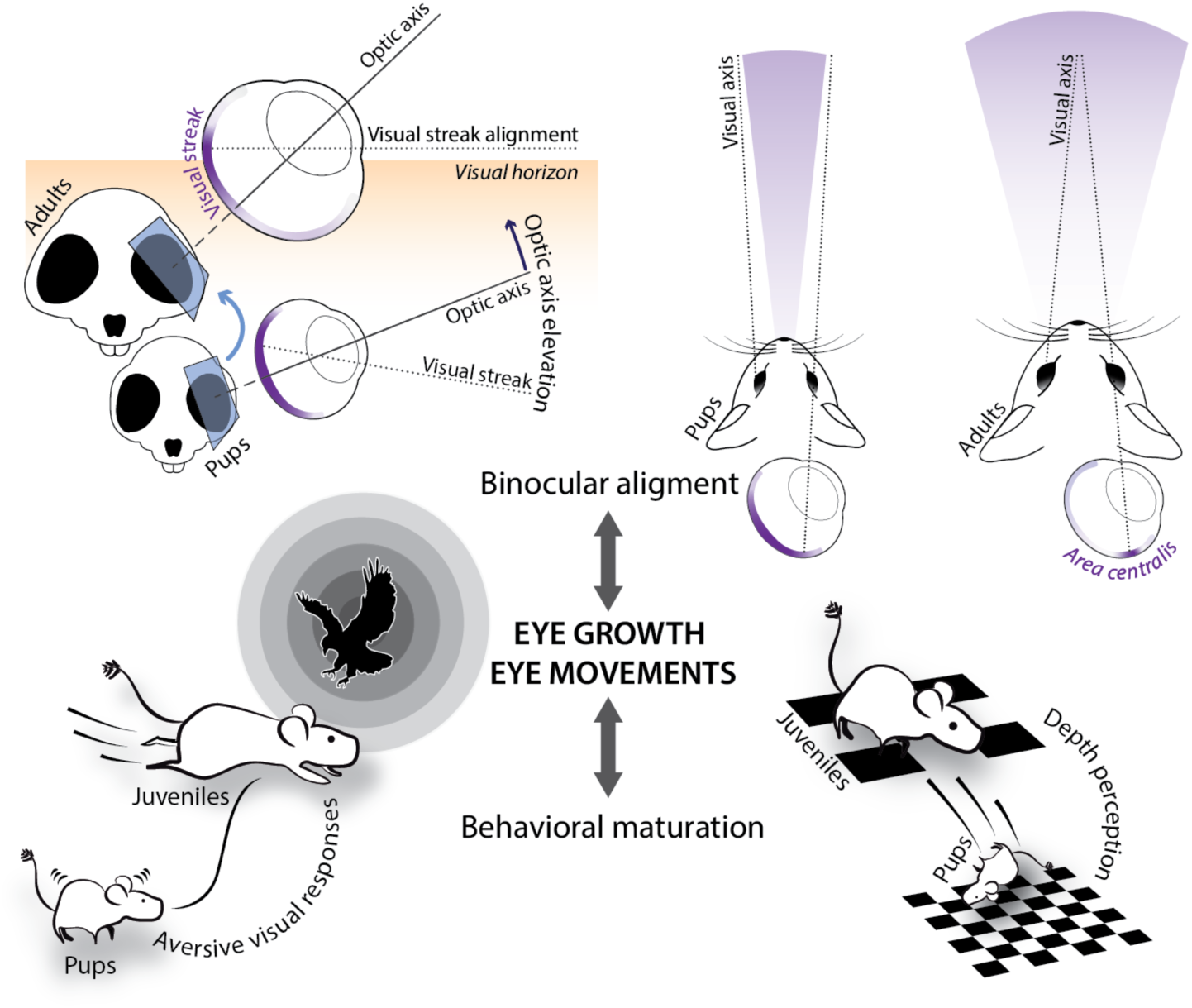
Summary schematics of the *Octodon degus* postnatal binocular ontogeny. Eye growth and eye movements are proposed as potential drivers of the postnatal integration of multiple binocular traits at different levels of organization.

Consistent with these anatomical changes, our behavioral tests show that binocular visually guided behaviors emerge postnatally, along with the maturation of the binocular visual field and retinal specialization. The visual cliff tests revealed a sharp emergence of depth discrimination between P9 and P15 in degus, which is at least eight days after the onset of visually guided locomotion.

Escape responses also show an ontogeny consistent with the emergence of binocular perception. Recent studies have reported that full escape probability develops after the third postnatal week in mice^19,20^. Here, we replicate and expand these results by showing that the development of escape responses follows different trajectories depending on the portion of the visual field being stimulated. The resulting asymmetry in behavior is coherent and contingent with the asymmetric expansion of the binocular field over the animal’s head. In adult rodents, escape initiation depends on the activation of the dorsal periaqueductal grey (dPAG) by medial SC neurons^68,69^, where the upper binocular field is represented^27,47–49^. A weak synaptic connection between medial SC and dPAG neurons imposes a threshold requiring high medial SC neuronal firing to elicit escape^68^. We suspect that binocular summation improves as the binocular field and ipsilateral retinal connections expand, resulting in enhanced activation of medial SC neurons. The sudden increase in escape at P15 could, therefore, reflect the threshold nature of the dPAG activation in response to visual stimulation.

Decades of research on the neural development of binocular vision have emphasized the role of molecular guidance^70,71^, spontaneous activity, neural plasticity and visual experience^72,73^ in shaping the connectivity and functional binocular responses in the brain, especially in V1 cortical neurons^18,74,75^. These processes are prominent in mice, where, for instance, unbalanced binocular inputs produce robust ocular dominance shifts in V1 and SC neurons^76,77^. Additionally, an initial mismatch in orientation preference for inputs from each eye in V1 binocular neurons is progressively corrected by visual experience^78–80^. Taken together, our results in degus indicate that binocular vision emerges during development through the coordinated alignment of multiple anatomical and functional components of the visual system. This process involves the integration of craniofacial morphology, retinal specialization, central visual pathways, and visually guided behavior within an active sensory context. These findings underscore the importance of developmental timing and cross-level interactions when studying the maturation of complex sensory functions. An integrative framework such as this is essential for understanding how sensory systems become functionally organized during ontogeny.

## Materials and Methods

### Animals

*Octodon degus* is a hystricomorph rodent endemic to central Chile. Degus were housed in metal cages (50 x 40 x 35 cm) with access to a running wheel and a refuge, in a climate-controlled room under a 12:12 photoperiod. They were fed a diet of rodent chow Prolab RMH 3000 (Purina LabDiet, Inc.), Mazuri chinchilla diet (Mazuri Exotic Animal Nutrition), alfalfa, and sunflower seeds, with free access to water. For reproduction, animals were kept in groups of three (one male and two females). Degus reproduce once or twice per year, and their gestation period lasts ∼3 months. All animal procedures were approved by the Institutional Committee for the Care and Use of Animals (CICUA) of the University of Chile.

### Visual field measurements

Animals were anesthetized in an induction chamber with 1.5–2% isoflurane in medical oxygen (100 mL/kg·min). Once unresponsive, diazepam (5 mg/kg, i.p.) was administered, followed by urethane (1 g/kg, i.p.) 15 min later. To suppress ocular movements during measurements, the extraocular muscles were infiltrated with a mixture of hyaluronidase (4%), lidocaine (1%), and DMSO (1%). Ophthalmoscopic procedures followed previously described protocols for adult degus and other mammalian species^7,28,30^. Animals were mounted in a stereotaxic head holder at the center of a campimeter, with the palpebral fissures aligned to the horizontal axis of the apparatus. Pupils were dilated with 2 μL of 1% tropicamide per eye, and eyelids were gently retracted with medical tape. Body temperature was maintained with a heating pad, and corneas were kept hydrated with sterile 0.9% NaCl. This step is critical to prevent ocular ischemia and lens opacification, which would preclude visualization of the retinal reflection. Eyes were examined with an ophthalmoscope. For each eye, monocular visual fields were delineated by measuring the borders of the retinal reflection across visual angles in 10 deg steps, and the optic axis was determined by aligning the corneal reflex with the center of the pupil. This was a terminal procedure; animals were euthanized for sample collection as described below. Visual fields were reconstructed in MATLAB (MathWorks) by projecting each monocular border onto a sphere representing the animal’s visual surround. Statistical tests were performed in R to evaluate the effect of age on binocular overlap (ANOVA), optic axis orientation (Kruskal-Wallis), and monocular field extension (Kruskal-Wallis), followed by post-hoc tests (t-tests, and Wilcoxon rank-sum tests) to compare differences across stages.

### Behavioral tests

#### Visual Cliff

We evaluated the performance of degus at P5, P9, P15 and adults (one trial per animal at only one of the stages). The experimental setup is a modification of the Gibson and Walk *Visual cliff* experiment^51^, and consisted of a glass arena (102 cm length x 40 cm width x 41 cm high) placed on a 92-cm-high platform with a checkerboard pattern. The arena extended 47 cm beyond a platform to simulate the “cliff”. The floor beneath the overhanging portion of the arena consisted of a checkerboard pattern with smaller square size than the platform pattern (see figure 5a). The walls were black, except for the frontal wall, which allowed the animal to have visual access to the outside. In addition, the back of the arena included a division with an opening to provide shelter to the animals and allow them to freely explore the arena.

We placed the animals in the shelter area and recorded them from above. Analyses were performed manually and restricted to the first encounter of the animals with the border of the cliff where the animals displayed two types of behaviors: either they kept moving forward without apparent disruption, scored as a crossing response, or they stop at the cliff border and after a few seconds withdrew to the opposite direction, scored as a non-crossing response. Fisher’s exact test was performed in R to determine if the performance on the cliff arena was associated with the developmental stage. Based on the cumulative scores for the animals in each stage tested, we elaborated a cliff Avoidance Index *(CAI) = (NC-NS)/(NC+NS)*, where NC is the number of animals that crossed the cliff and NS is the number of animals that stopped before the cliff. An Avoidance Index close to 1 indicates a preference for stopping and not crossing the cliff, while an index closer to −1 indicates a tendency to cross the cliff border at the corresponding stage.

#### Looming response

Reproducing the set up described in our previous study in adult degus^47^, we presented aerial or lateral looming stimuli and recorded the behavioral responses. Animals were placed in an arena 48 cm wide x 35 cm deep x 30 cm high, equipped with 2 monitors, one placed in one of the walls of the arena for frontal stimulation and the other on the top for the overhead looming display. For each trial, a stimulus was presented in one of these monitors when the animal was freely moving, with both monitors in the field of view. Stimuli were generated using the open-source software Psychopy in Python76. Degus movements were video recorded with a camera, using VCL Media Player (30 frames/s) and tracked with the software DeepLabCut^81^.

Analyses were made using a Python custom-made routine. Fisher’s exact test was performed using software R to determine differences in response to the location of looming stimulation across different stages. Instantaneous speed was calculated as the trajectory covered by the animal between two consecutive frames, pixels/frame were converted into cm/s. The looming stimulus consisted of a 2° black disk rapidly widening to 50° in a 250-ms lapse. Stimuli were presented randomly to each individual. The degus were habituated to the box for 15 min. the day before the first trial. Each animal was stimulated only once with a given stimulus in single trials at least three days apart.

#### Intraocular injections of neural tracers

We performed intraocular injections of the cholerotoxin B subunit neural tracer to visualize the retinal projections to the brain. Pups and juveniles were injected 24 h before perfusion at stages P4, P8, P14, and P29. Adult projection data were quantified from material from a previous study from our laboratory^7^.

After sedation with isoflurane as described above, diazepam (5 mg/kg, i.p.) was administered and animals remained in the isoflurane chamber for a further 5 min. Animals were then transferred to a heating pad and anesthesia was maintained via a modified silicone nose mask with 2% isoflurane at 50 mL/kg·min, providing deep, stable anesthesia with rapid recovery. A Hamilton syringe was used to deliver 6 - 10 μL of 0.5% cholera toxin subunit B (CTB; List Biological Laboratories) in PBS with 1% DMSO into the vitreous chamber of the left eye. Proparacaine 0.5% was applied topically at the injection site, and carprofen (5 mg/kg, s.c.) was given as an analgesic. Pups were returned to their littermates and after 24 h, animals were euthanized and perfused as described below.

#### Tissue collection

At the end of all procedures, animals were euthanized to obtain skulls, retinas, and brains, by placing them in an induction chamber infused with 1.5–2% isoflurane in medical oxygen, delivered at a rate of 100 ml/kg·min until unresponsive, then given an intraperitoneal overdose of ketamine/xylazine. After animals ceased breathing, they were intracardially perfused with 100 mL of 0.1 M phosphate saline solution (PBS), followed by 100 mL of 4% paraformaldehyde (PFA) in PBS. Brains were removed through a craniotomy at the skull base to preserve the orbits and cranial vault for subsequent measurements of orbital convergence.

Following fixation, marks were made with a hot needle onto the palpebral fissures and dorsal sclera of the eyeballs for orientation. The eyes were enucleated and postfixed in 4% PFA for 1 hour and then washed in 0.1 M PBS. Retinas were carefully dissected from the underlying pigmented layer, and the optic nerves were severed just beneath the optic disc.

### Tissue preparation and imaging

#### Skull CT-scan

After all soft tissue was removed, the skulls were imaged using X-ray micro-Computed Tomography Scanner (SkyScan 1278) at a resolution of 51.48μm. Volumes were reconstructed using the 3DSlicer open-source software. Three anatomical landmarks were manually placed to define the sagittal plane and three per orbit to define orbital planes, following the protocol described by Heesy^9^. These coordinates enabled calculation of the dihedral angle between each orbital plane and the sagittal plane. Non-parametric statistical tests were performed in R to evaluate the effect of age on orbit convergence (Kruskal–Wallis), followed by post-hoc tests (Wilcoxon rank-sum tests) to compare differences across stages and a Pearson’s correlation to evaluate the covariation between orbital convergence and binocular field during development.

#### Retinal wholemount immunofluorescence

For immunohistochemistry in floating whole retinas, we followed the protocol described by D’Souza et al^32^. Retinas were incubated in 0.5% Triton-X in PBS (0.5% PBST) for 15 min, followed by 15 min incubation in 50% acetone in dH_2_O for antigen retrieval. Blocking was performed with 4% Normal Horse Serum (NHS) and 0.5% Triton in PBS (30 min), followed by primary antibody (1:200, rabbit anti-RBPMS, Sigma) incubation in 3% NHS and 0.3% Triton at 4 °C for 5 days. After PBS washes, retinas were incubated with fluorescent anti-rabbit secondary antibody (1.5% NHS, 0.15% Triton in PBS) for 2 days at 4 °C in darkness. Finally, retinas were mounted on glass slides using radial cuts to flatten the tissue while maintaining the original dorsal orientation. Specimens were slowly air-dried in a humid dark chamber at 4 °C and cover slipped using FluorSave reagent (Millipore) as mounting medium.

Each retina was imaged using a confocal microscope with a 20X objective. Retinal ganglion cell quantification was then performed using StereoInvestigator software for assisted counting, 5% of the retinal surface was sampled using a 150 μm grid. Total cell number and retinal surface area were estimated.

Using the spatial data obtained in Stereoinvestigator, isodensity maps were generated for each retina using the script provided by Garza-Gisholt et al.^82^ on RStudio, applying the “Thin plate spline” function with a smoothing factor equivalent to two-thirds of the sampling sites. This approach enabled the construction of spatial density maps along with an estimation of peak cell density and lower cell density values to compute central-to-peripheral (C/P) ratios for each retina. An ANOVA was performed to determine statistical changes in the number of RGCs; non-parametric statistics (Kruskal-Wallis followed by Wilcoxon rank-sum tests) to examine changes in retinal area and visual streak coverage, and a linear regression to evaluate the ontogenetic correlation between retinal area and C/P ratios.

#### Brain tissue processing and analysis

Brains were processed for CTB visualization as previously described in the degus^47,83^. After their removal from the skull, brains were postfixed in 4% PFA in PBS for 24h and placed in a 30% sucrose solution until they sank. Using a freezing microtome, 40 μm-thick sections were obtained and collected into 4 series of sections. One series of each brain was then washed in PBS and incubated in hydrogen peroxide solution (3%) to quench endogen peroxide activity, prior to incubation in anti-CTB primary antibody solution (List Labs) diluted 1:20000 in a blocking solution containing 0.3% Triton X-100 and 3% normal horse serum overnight. Then, sections were washed in PBS and incubated in secondary anti-goat antibody (Jackson ImmunoResearch Labs) diluted 1:2000 in blocking solution for 2h at room temperature. Sections were then rewashed in PBS and incubated in avidin–biotin peroxidase complex solution (1: 750; ABC kit, Vector) for 2h. To reveal peroxidase binding, sections were washed and then incubated in diaminobenzidine and hydrogen peroxide for 5 - 10 min. After washing in PBS, sections were mounted in gelatin coated slides, dehydrated in a series of alcohols and xylene, and cover-slipped with mounting medium (Entellan, Merck Millipore).

Slides were photographed on a slide scanner (EasyScan, Motic) equipped with a 40x objective, and photos of the sections of each case were aligned using Fiji. Then, z-stacks were imported into a 3D image-analysis software (VG studio Max, Volume Graphics, Heidelberg, Germany) to estimate the volumes of CTB labeling in SC and dLGN ipsi- and contralateral to the injection. Non-parametric statistical tests were performed in R to evaluate global effects (Kruskal–Wallis) and monotonic changes across postnatal stages (Jonckheere–Terpstra).

## Acknowledgments

We thank Dr. Laura Fenlon, Dr. Rodrigo Suarez, Dr. Cristian Gutierrez, Dr. Alfredo Kirkwood, and Dr. Alexander Vargas for their review and comments on the draft. We gratefully acknowledge Elisa Sentis and Solano Henriquez from El Rayo lab for their technical support, Lorena Saragoni (Unidad de Microscopía Avanzada, Departamento de Biología, Facultad de Ciencias, U. de Chile) and Daniela Poblete (Plataforma Experimental Bio-CT, Facultad de Odontología, U. de Chile) for their invaluable assistance with confocal microscopy and CT-scanning, and Francisca Silva for her help in editing of supplementary videos.

This work was supported by ANID grant Fondecyt Postdoctorado (Chile): 3220871(A.D.), ANID grant Fondecyt Regular: 1210169 (G.M.), ANID Millennium Nucleus Early Evolutionary Transitions of Mammals (EVOTEM): NCN2023_025 (M.F.), ANID grant Fondecyt Postdoctorado: 3220759 (C.M.) ANID grant Fondecyt Regular: 1250880 (J.M.).

## Author Contributions

A.D., M.R.-F., N.I.M., C.M., L.L.-J., J.M., M.F. and G.M. designed research; A.D., M.R.-F., N.I.M., L.L.-J., T.V.-Z., and M.F. performed research; A.D., M.R.-F., N.I.M., C.M., L.L.-J., and M.F analyzed data; A.D., M.F. and G.M. wrote the paper.

