## supplementary table for "Binocular vision emerges from the coordinated development of orbit convergence, eye orientation, and high-acuity retinal specializations"

**This PDF file includes:**

Tables S1 to S3
Legends for Movies S1 to S2

**Other supporting materials for this manuscript include the following:**

Movies S1 to S2

**Tables**

**Table S1.** Estimated monocular visual field, binocular visual field, and dihedral angle between the orbit and the midsagittal plane in *Octodon degus* at different postnatal stages.

| | binocular overlap (°) | | monocular field (°) | | optic axis elevation (°) | | dihedral angle (°) | |
|---|---|---|---|---|---|---|---|---|
| Stage | Average | SD | Average | SD | Average | SD | Average | SD |
| P5 | 25.5 | 5.1 | 166.6 | 15.3 | 36.3 | 12.5 | 20.1 | 2.7 |
| P9 | 34.2 | 7.0 | 177.9 | 7.8 | 44.6 | 10.8 | 22.2 | 2.9 |
| P15 | 40.4 | 8.9 | 174.6 | 9.0 | 55.1 | 8.5 | 27.9 | 1.6 |
| P30 | 54.2 | 6.8 | 183.3 | 7.3 | 56.7 | 7.7 | 39.4 | 1.4 |
| Adult | 59.5 | 11.4 | 186.3 | 7.2 | 63.0 | 5.2 | 43.4 | 2.3 |

**Table S2.** Estimated retinal area, RGC number, RGC density, Peak RGC density, and density ratio between central and peripheral retina in *Octodon degus* at different postnatal stages.

| | Retinal area (mm2) | | RGC number | | mean RGC density (cell/mm2) | | peak RGC density | | C/P ratio | |
|---|---|---|---|---|---|---|---|---|---|---|
| Stage | Average | SD | Average | SD | Average | SD | Average | SD | Average | SD |
| P1 | 36.6 | 2.7 | 163840.3 | 3710.9 | 4493.6 | 390.7 | 7224.6 | 1245.6 | 9.6 | 3.3 |
| P5 | 42.7 | 7.5 | 184295.3 | 13524.6 | 4468.4 | 1083.4 | 7122.0 | 1494.4 | 8.9 | 3.6 |
| P8-9 | 45.3 | 6.8 | 173568.4 | 8836.5 | 3937.4 | 568.9 | 6925.3 | 747.7 | 11.0 | 5.4 |
| P15 | 54.0 | 5.1 | 180828.2 | 14986.5 | 3414.1 | 489.3 | 6129.3 | 1012.3 | 9.8 | 1.8 |
| P31 | 64.2 | 5.4 | 174575.0 | 14388.5 | 2680.8 | 167.0 | 5369.0 | 531.9 | 15.1 | 6.6 |
| Adult | 96.5 | 4.8 | 178693.7 | 11962.4 | 1855.2 | 148.5 | 4701.8 | 386.5 | 17.0 | 3.0 |

**Table S3.** Estimated volume of dLGN and SC labelling after CTB injection in one eye at different postnatal stages in *Octodon degus*.

| | Ipsilateral dLGN labelling volume ($mm^3$) | | Ipsilateral SC labelling volume ($mm^3$) | | Contralateral dLGN labelling volume ($mm^3$) | | Contralateral SC labelling volume ($mm^3$) | |
|---|---|---|---|---|---|---|---|---|
| Stage | Average | SD | Average | SD | Average | SD | Average | SD |
| P5 | 0.0050 | 0.0046 | 0.0007 | 0.0001 | 0.7541 | 0.1614 | 2.7150 | 0.4228 |
| P9 | 0.0101 | 0.0100 | 0.0047 | 0.0039 | 0.7493 | 0.1004 | 2.7248 | 0.4379 |
| P15 | 0.0116 | 0.0047 | 0.0017 | 0.0009 | 1.0542 | 0.0814 | 3.3346 | 0.8432 |
| P30 | 0.0199 | 0.0086 | 0.0073 | 0.0065 | 1.6231 | 0.1584 | 3.7811 | 0.5289 |
| Adult | 0.0112 | 0.0158 | 0.0078 | 0.0039 | 1.4670 | 0.2044 | 3.2867 | 0.7946 |

**Movie S1 (separate file). Postnatal emergence of depth perception.** This movie shows two representative examples of degus performing the visual cliff test at stages P9 and P15, illustrating the developmental shift in depth-related responses. At P9, the individual explores the arena without reacting to the cliff. By contrast, at P15, the individual pauses at the edge and avoids the drop, reflecting the emergence of depth discrimination.

**Movie S2 (separate file). Postnatal maturation of escape responses.** This movie illustrates how asymmetric changes in aversive responses to looming stimulation align with the maturation of the binocular visual field. The video shows individuals at stages P9 and P15 to highlight the developmental shift in behavior. At P9, the individual exhibits freezing responses to looming stimuli presented either in the upper or frontal regions of the visual field. By contrast, at P15, the individual freezes only to frontal looming stimuli, while upper-field stimulation elicits a rapid escape reaction.
